# Autism-associated Shank3 is essential for homeostatic plasticity and neuronal circuit stability

**DOI:** 10.1101/365445

**Authors:** Vedakumar Tatavarty, Alejandro Torrado Pacheco, Heather Lin, Nathaniel J. Miska, Keith B. Hengen, Florence F. Wagner, Gina G. Turrigiano

**Author notes:** Current address: Emory University Medical School, Atlanta, GA,30322. Current address: Dept of Biology, Washington University, St. Louis, MO 63130.

## Abstract

Mutations in Shank3 are strongly associated with autism spectrum disorders and circuit disfunction, but a unified view of how Shank3 loss disrupts circuit function and excitability is lacking. Stabilizing, homeostatic forms of synaptic and intrinsic plasticity are critical for preventing circuit hyper- or hypo-excitability, leading us to ask whether Shank3 loss perturbs circuits by disrupting homeostatic plasticity. We show that Shank3 loss abolishes synaptic and intrinsic homeostatic plasticity, which can be rescued by lithium(Li), a drug with therapeutic potential in human Shankopathies. Further, Shank3 loss *in vivo* severely compromises the ability of visual cortical circuits to recover from perturbations to sensory drive. Our findings suggest that the loss of homeostatic compensation is a critical endophenotype that can explain a range of circuit disfunctions in Shankopathies.

## INTRODUCTION

Mutations in the multidomain scaffold protein Shank3 are associated with autism spectrum disorders (ASDs), Phelan-McDermid syndrome, and intellectual disability (Betancur and Buxbaum, 2013). Animal models of Shank3 loss have uncovered a number of circuit defects in a wide range of brain regions that might contribute to the behavioral phenotypes of these Shankopathies (Betancur and Buxbaum, 2013; Wilson et al., 2003; Zhou et al., 2016), but a unified view of how haploinsufficiency of Shank3 disrupts circuit function is lacking. A common feature of Shankopathies (as well as other monogenic ASDs) is the development of hyper- or hypoexcitability within specific CNS circuits (Monteiro and Feng, 2017; Nelson and Valakh, 2015); because homeostatic plasticity normally serves to prevent such circuit imbalances from developing, this raises the intriguing possibility that defects in homeostatic plasticity represent a common endophenotype of monogenic ASDs (Blackman et al., 2012; Dani et al., 2005; Nelson and Valakh, 2015). While loss of Shank3 has been reported to modulate baseline synaptic transmission and intrinsic excitability in some brain regions and neuron types (Jiang and Ehlers, 2013; Peca et al., 2011; Yi et al., 2016), a role for Shank3 in homeostatic plasticity has not been explored.

Maintenance of circuit stability in the neocortex is thought to be achieved through a set of homeostatic plasticity mechanisms, including synaptic scaling (Turrigiano et al., 1998) and intrinsic homeostatic plasticity(Desai et al., 1999), that cooperate to stabilize neuronal firing (Hengen et al., 2013; Hengen et al., 2016; Lambo and Turrigiano, 2013). Synaptic scaling bidirectionally scales all excitatory postsynaptic strengths up or down in the right direction to compensate for prolonged perturbations in firing (Turrigiano et al., 1998), while intrinsic homeostatic plasticity adjusts the input-output function of neurons by regulating the balance of inward and outward voltage-dependent conductances (Desai et al., 1999; Joseph and Turrigiano, 2017; Lambo and Turrigiano, 2013). Recent experiments in the rodent primary visual cortex (V1) indicate that perturbing firing through monocular eyelid suture (MD) induces both types of homeostatic plasticity (Kaneko et al., 2008; Keck et al., 2013; Lambo and Turrigiano, 2013; Mrsic-Flogel et al., 2007), which then cooperate to restore neuronal firing back to an individual firing rate set point (Hengen et al., 2013; Hengen et al., 2016), a process referred to as firing rate homeostasis (FRH). The possibility that defects in FRH might contribute to circuit instability has not been tested in any ASD model.

Our data demonstrate that a single genetic manipulation, the loss of Shank3, results in the simultaneous loss of both synaptic and intrinsic homeostatic plasticity in neocortical pyramidal neurons. Surprisingly both these deficits can be rescued *in vitro* by treatment with Li or by the direct pharmacological inhibition of a Li target, Glycogen Synthase Kinase 3 (GSK3). Finally, multielectrode array (MEA) recordings in awake freely behaving Shank3 knockout mice demonstrate that these homeostatic defects result in a dramatic loss of FRH. These results support a causal relationship between loss of homeostatic mechanisms and deficits in FRH. Furthermore, they suggest that loss of homeostasis may contribute to circuit disfunctions in ASDs.

## RESULTS

### Shank3 is required for synaptic scaling

Shank3 is a multidomain scaffold protein enriched at postsynaptic sites where it interacts with a number of synaptic scaling effector proteins (Monteiro and Feng, 2017). We found that synaptic scaling paradigms bidirectionally regulated the synaptic localization of Shank3: blocking spiking with tetrodotoxin (TTX) for 24 hr recruited both synaptic AMPA-type glutamate receptors (GluA2) and Shank3 to excitatory synapses (Fig 1A,B), while enhancing spiking with picrotoxin (PTX) for 24 hr reduced both GluA2 and Shank3 synaptic localization (Fig. 1B). To determine if a normal complement of Shank3 is necessary for synaptic scaling, we used RNAi to induce an acute and sparse knockdown (KD) of Shank3 (Verpelli et al., 2011) to ∼50% of control levels in individual pyramidal neurons (Fig. S1A,B). Shank3 KD (SH) had no significant impact on the basal amplitude of miniature excitatory postsynaptic currents (mEPSCs) compared to empty-vector (EV) controls (Fig. 1C-F), or on mEPSC frequency or excitatory synapse density (Fig. S2C,D). In contrast, Shank3 KD completely blocked both TTX-induced synaptic scaling up (Fig. 1C-F), and PTX-induced synaptic scaling down (Fig. 1G-H) of mEPSC amplitude, as well as the changes in GluA2 synaptic accumulation that drive synaptic scaling (Fig. S2A, B, E and F). Synaptic scaling could be rescued by co-expression of an RNAi-resistant form of Shank3 (Fig. 1F, RES), ruling out off-target effects of the shRNA. Thus, a 50% loss of Shank3 (analogous to haploinsufficiency in human Shankopathies) is sufficient to completely abolish bidirectional synaptic scaling, and this requirement is cell-autonomous.

**Fig 1.**
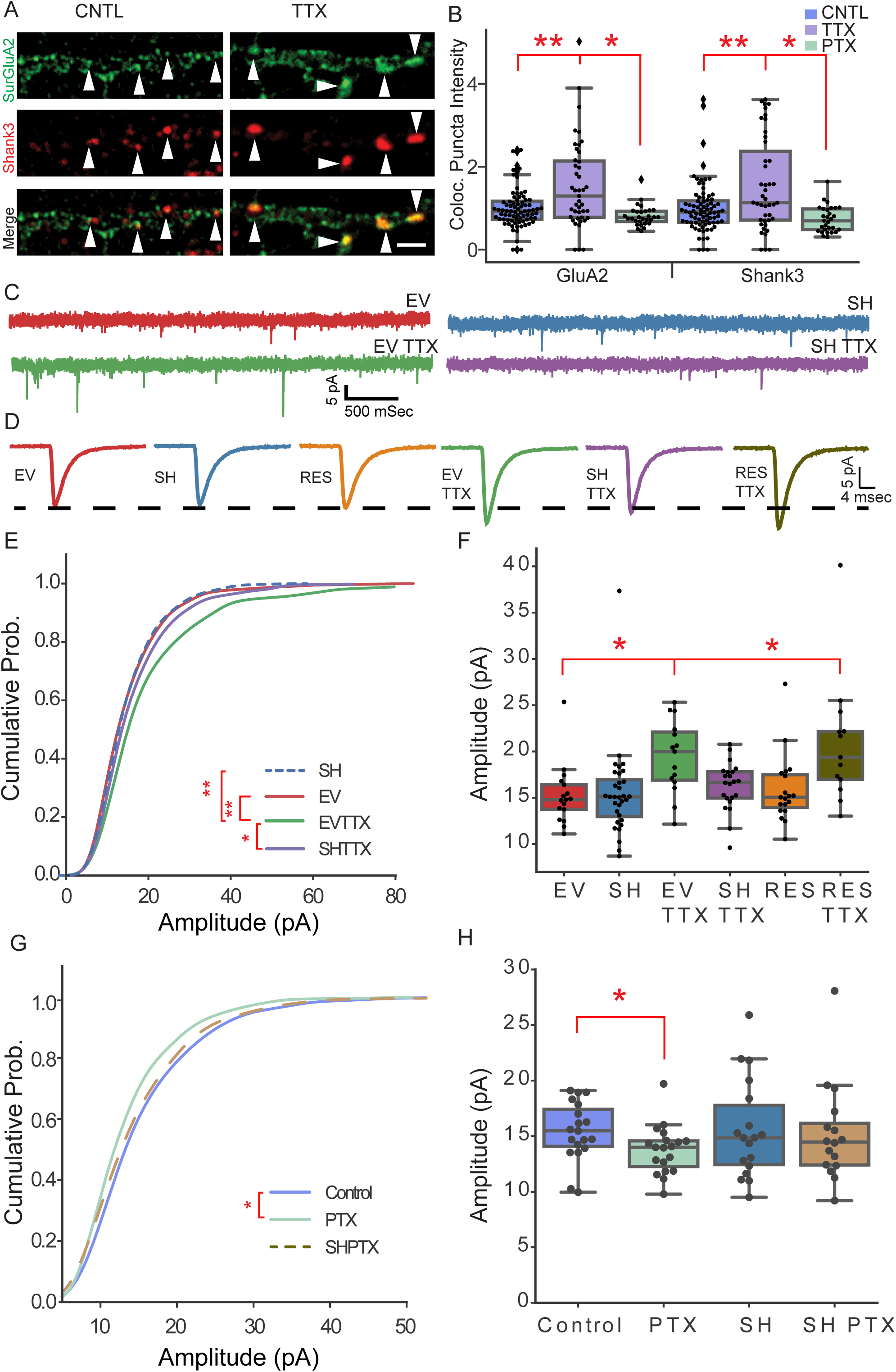
Block of bidirectional synaptic scaling by Shank3 loss. (A) Representative images of control (left) and TTX-treated (right) dendrites, immunolabled for surface GluA2 (green) and Shank3 (red). Arrowheads indicate example colocalized puncta. Scale bar 2 μm. (B) Total intensity of GluA2 (left) and Shank3 (right) signal at colocalized sites after 24 hrs in TTX or PTX (*n,* Control = 80, TTX = 43, PTX = 29 neurons);ANOVA followed by Bonferroni correction (BC). (C) Example raw traces and (D) average mEPSCs waveforms for indicated conditions. (E) Cumulative histograms of mEPSC amplitudes; Kruskal-Wallis (KW) test with BC. (F) mean mEPSC amplitude, (*n,* EV = 17, EV TTX = 15, SH = 32, SH TTX = 22, RES = 19, RES TTX = 13); One-way ANOVA followed by BC. (G) Cumulative histograms of mEPSCs; KW test with BC. (H) Mean EPSC amplitude (*n,* Control = 19 PTX = 19, SH = 17, SH PTX = 18); two-tailed Student’s t-test. SH PTX not different from SH, p>0.05. Here and below, Box plots: box depicts first and third quartiles of the data and whiskers show remainder of the distribution. Line depicts median. **indicates p<0.01, * indicates p<0.05; p>0.5 considered N.S. Actual p values for each comparison are given in suppl. Table S1.

### Synaptic scaling blockade caused by Shank3 KD can be relieved by Li treatment

Li has wide efficacy to treat many neurological disorders, but the mechanism of action is debated (Pisanu et al., 2016). Several recent case studies suggest that Li has therapeutic potential for treating human Shankopathies (Darville et al., 2016; Egger et al., 2017; Serret et al., 2015), leading us to wonder whether Li might rescue the synaptic scaling defect induced by Shank3 loss. After 24 hr incubation with a therapeutic dose (1 mM) of LiCl, Shank3 KD neurons were once again able to fully express synaptic scaling (Fig 2A-C). Li did not exert these effects through a simple global enhancement of synaptic scaling, as Li treatment had no impact on the magnitude of scaling in control neurons with a normal Shank3 complement (Fig. S3A,B). Further, although prolonged Li treatment (2 mM) is reported to reduce mEPSC amplitude in hippocampal neurons (Gideons et al., 2017), here Li had no significant impact on basal postsynaptic (Fig. S4 B,C) or neuronal (Fig. S3) properties. A major target of Li is GSK3; Li acts as a direct and indirect inhibitor of GSK3 through the inhibitory phosphorylation of Ser9/21 (Freland and Beaulieu, 2012). We therefore tested a highly specific GSK3 inhibitor BRD0320 (GSK3i), that blocks GSK3 alpha and beta but not other closely related kinases (Wagner et al., 2016), for its ability to rescue synaptic scaling. Like Li, BRD0320 was able to rescue synaptic scaling in Shank3 KD neurons (Fig. 2D), suggesting that the GSK3 pathway is dysregulated by Shank3 KD in neocortical pyramidal neurons, and contributes to the defect in synaptic scaling.

**Fig 2.**
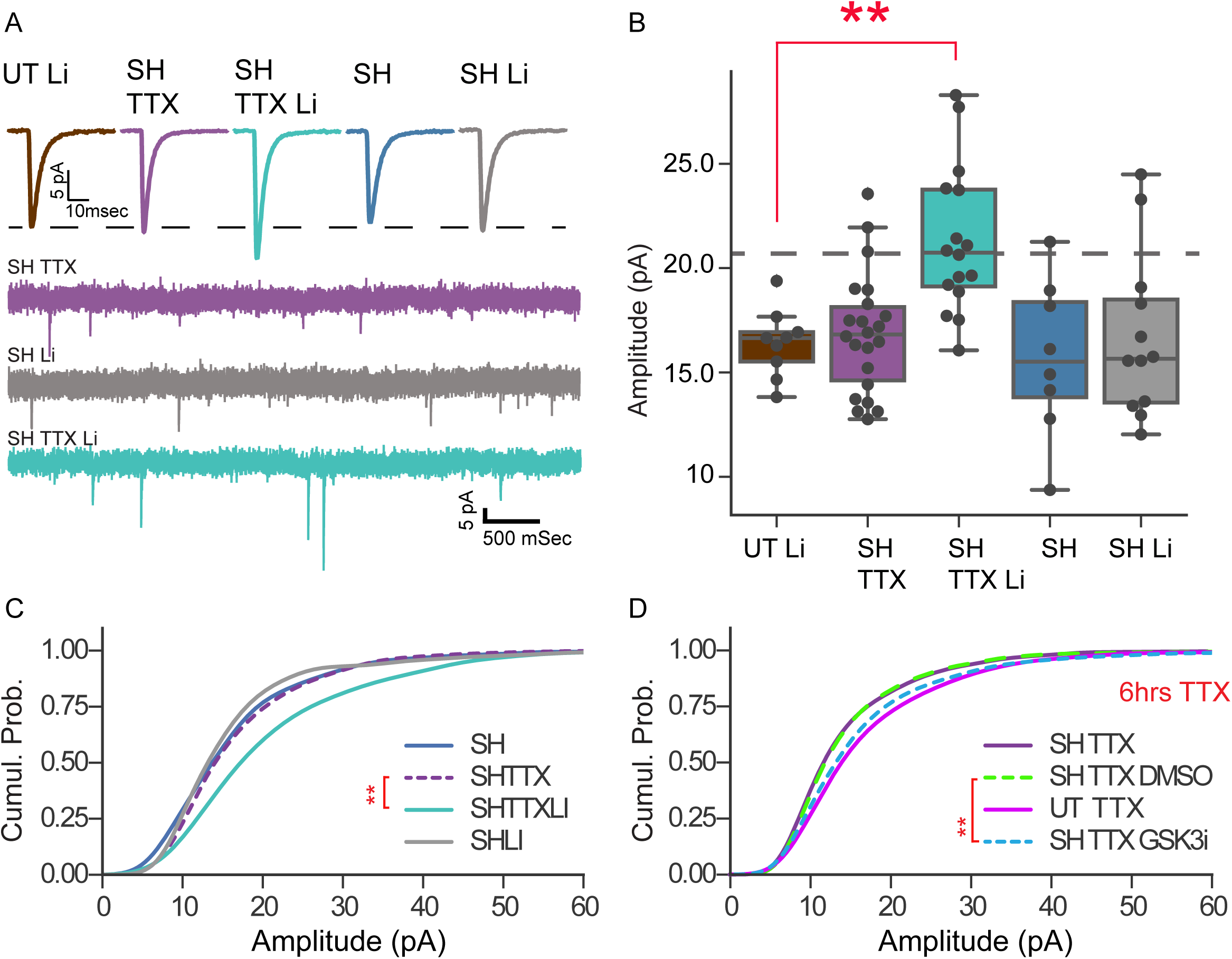
LiCl treatment rescues synaptic scaling. (**A**) Average mEPSC traces (top) and example raw traces (below) from control or knockdown neurons treated with Li or Li + TTX for 24 hrs. (**B**) Mean mEPSC amplitudes (*n*, Li = 9, SH TTX = 22, SH TTX Li = 16, SH = 8, SH Li = 12); One-way ANOVA followed by BC. (**C**) Cumulative histograms of mEPSCs for conditions in (B); KW test with BC. (**D**) Cumulative histograms from neurons treated for 6 hrs with TTX + DMSO (vehicle) or TTX + 5 μM GSK3 inhibitor (GSK3i) (*n*, UT TTX = 30, SH TTX = 25, SH TTX DMSO = 31, SH TTX GSK3i = 30. SH TTX GSK3i vs SH TTX DMSO (**); KW test, with BC.

### Shank3 is required for homeostatic regulation of intrinsic excitability

Shank3 interacts with ion channels as well as synaptic scaffold proteins (Yi et al., 2016), but the possibility that intrinsic homeostatic plasticity is impaired in shankopathies (or any other ASD model) has not been explored. To determine whether Shank3 is essential for intrinsic homeostatic plasticity as well as synaptic scaling, we knocked down Shank3 as before, blocked spiking for 24 hr, and then probed intrinsic excitability by generating FI curves in the presence of synaptic blockers (Fig. 3). After TTX treatment the same current injection evoked more action potentials in EV neurons (Fig. 3A), and the entire FI curve shifted upward/leftward (Fig. 3B) as expected (Desai et al., 1999). Interestingly, although Shank3 KO reduces a hyperpolarization-activated cation current (I_h_) and thus increases input resistant in hippocampal pyramidal neurons (Yi et al., 2016), Shank3 KD had no impact on passive neuronal properties (Fig. 3D,F) or on basal intrinsic excitability (Fig. 3B,C) in these neocortical pyramidal neurons; similarly, input resistance was not altered in L2/3 pyramidal neurons from Shank3 KO mice relative to WT littermates (Fig. 4.C,D., S5). These data are consistent with the existence of cell type-specific differences in Shank3-ion channel interactions (Zhu et al., 2018). Although baseline excitability was largely unaffected by Shank3 KD, intrinsic homeostatic plasticity was completely absent in Shank3 KD neurons (Fig 3B,C). Strikingly, we found that Li treatment could rescue intrinsic homeostatic plasticity (Fig. 3G,H) as well as synaptic scaling.

**Fig 3.**
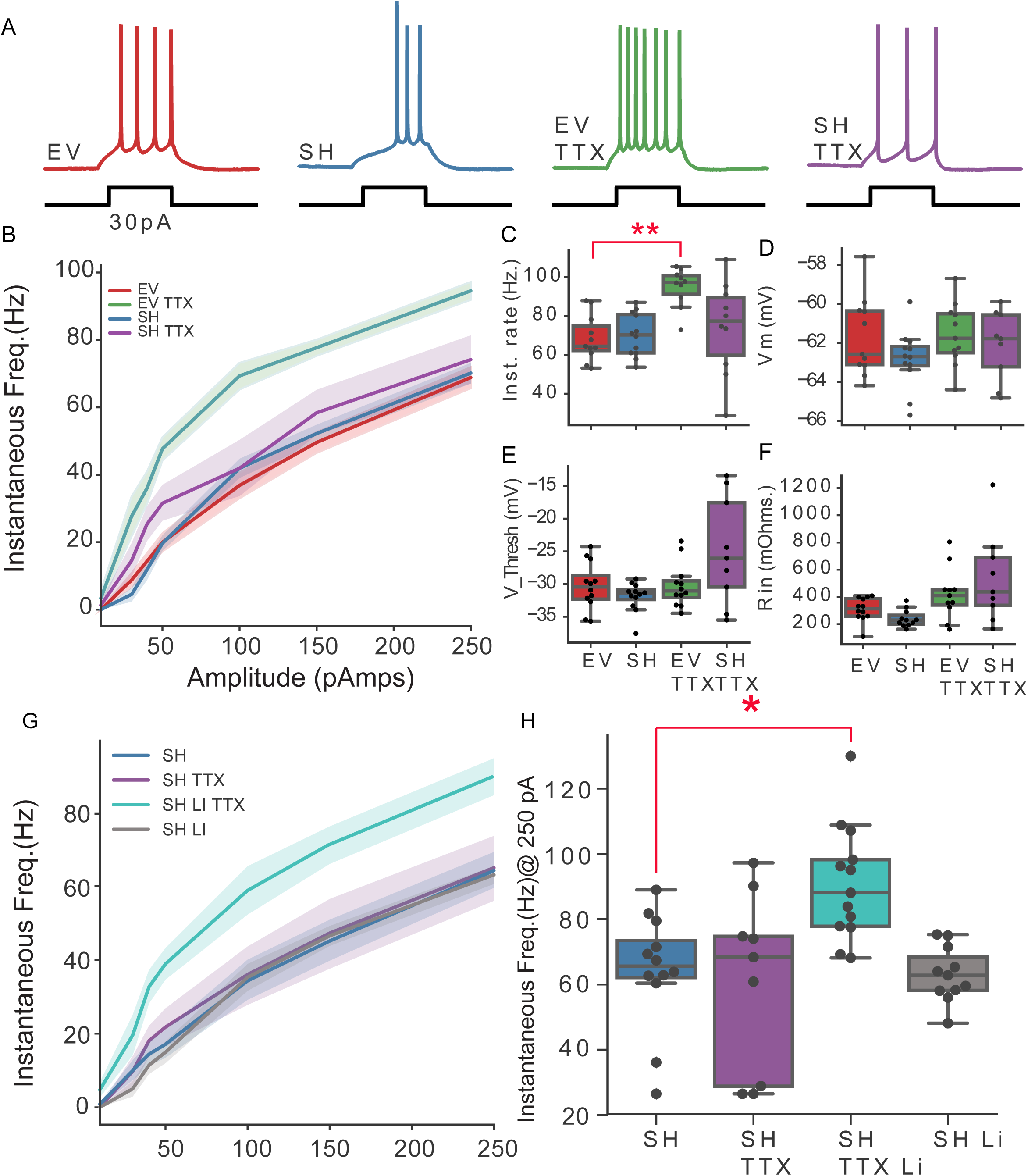
Shank3 loss blocks, and Li rescues, intrinsic homeostatic plasticity. (**A**) Example firing elicited with DC current injection from neurons in indicated conditions. (**B**) Average *f*-I curves for indicated conditions. Shaded area around curves denotes standard error. (**C**) Average firing rates for indicated conditions during highest current injection (250 pA). (*n*, EV = 12, EV TTX = 12, SH = 11, SH TTX = 10); One-way ANOVA with BC. (**D-F**) Passive neuronal properties across conditions; VThresh = voltage threshold for action potential firing. (**G**) Li rescue: *f*-*I* curves for indicated conditions. (**H**). Average firing rates from conditions in (G) at highest current injection (250 pA). (*n*, SH = 13, SH TTX = 11, SH TTX Li = 13, SH Li = 11); One-way ANOVA with BC.

**Fig 4.**
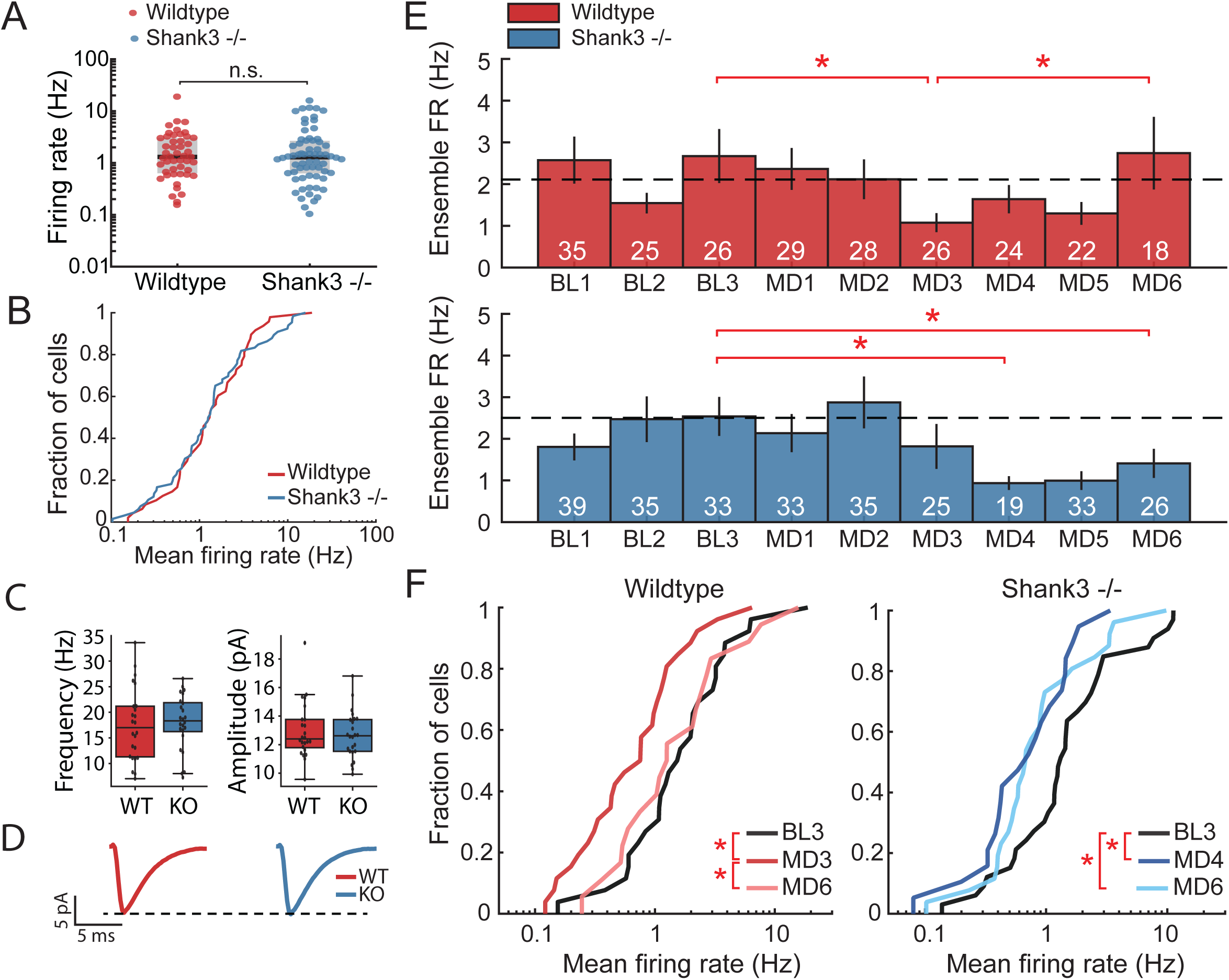
Shank3 is required for firing rate homeostasis *in vivo*. **(A)** Spontaneous mean firing rates in V1 of freely behaving mice at baseline (BL2 and BL3) for WT (red) and Shank3 KO (blue) (N.S.), Wilcoxon rank-sum test. **(B)** Cumulative distributions of baseline firing rates; Anderson-Darling test. **(C)** Frequency (left) and amplitude (right) of V1 L2/3 pyramidal neuron mEPSCs (*n*, WT and KO = 24; two-sample t-test (N.S). **(D)** Average mEPSC waveforms from WT or KO mice. **(E)** Ensemble average firing rates (Hz) in V1 of freely behaving mice during MD; KW test followed by pairwise Wilcoxon rank-sum tests with Benjamini-Hochberg FDR procedure (BH-FDR). Bars show mean ± s.e.m. White numbers within bars indicate number of neurons. **(F)** Cumulative distributions of mean firing rates comparing baseline, early, and late MD; Anderson-Darling test followed by BH-FDR.

### Shank3 knockout mice have deficits in firing rate homeostasis

A major function of synaptic scaling and intrinsic homeostatic plasticity *in vivo* is to stabilize neuronal firing over long timescales through firing rate homeostasis (Hengen et al., 2013; Turrigiano, 2017). Given that loss of Shank3 disrupts both of these major forms of homeostatic plasticity, we wondered whether firing rate homeostasis *in vivo* would also be disrupted. To test this, we implanted critical-period Shank3 KO (Peca et al., 2011) or littermate WT mice with multielectrode arrays in V1, performed MD, and followed changes in firing rates over time in freely behaving animals. In both WT and KO animals the distributions of baseline firing rates were wide as expected (Darville et al., 2016; Peixoto et al., 2016), and were similar between WT and KO (Fig. 4A). MEPSC properties were also similar between WT and KO L2/3 pyramidal neurons recorded in acute slices (Fig. 4C,D). In freely behaving WT mice MD resulted in biphasic changes in the firing of V1 neurons as expected (Hengen et al., 2013; Hengen et al., 2016): first a drop in mean firing rates that reached a minimum after 3d of MD (due to suppressive Hebbian mechanisms induced by blurred vision through the closed eye (Smith et al., 2009), followed by a homeostatic restoration of firing rates back to baseline values (Fig. 4E,F). However, the behavior of neurons in the KO was strikingly different: mean firing rates dropped as for WT animals (although more slowly, reaching a nadir after 4d of MD), but there was no recovery in firing rates over the ensuing days (Fig. 4E). In WT animals the entire distribution of firing rates initially shifted to the left, then recovered so by MD6 the distribution was statistically indistinguishable from baseline (Fig 4F left). In contrast, in Shank3 KO mice the distribution shifted to the left and there was little recovery by MD6 (Fig. 4F, right). These data indicate that loss of Shank3 severely compromises the ability of neocortical neurons to recover stable function after perturbations in drive; this normal recovery is either absent or dramatically slowed.

## DISCUSSION

Homeostatic compensation is widely considered essential for neurons and circuits to maintain (or restore) stable function during development and experience-dependent plasticity (Abbott and Nelson, 2000; Davis, 2013; Marder and Goaillard, 2006; Turrigiano and Nelson, 2004). Here we show that loss of the autism-associated multidomain scaffold protein Shank3 abolishes both major forms of homeostatic compensation in neocortical pyramidal neurons: synaptic scaling and intrinsic homeostatic plasticity. Further, we find that Shank3 loss prevents the normal homeostatic recovery of V1 firing rates during prolonged visual deprivation, indicating that Shank3 loss severely compromises circuit stability. These data suggest that a major sequela of Shank3 loss is the inability of neocortical circuits to compensate for the changes in synaptic drive that are a normal aspect of circuit development and critical period plasticity.

In humans, Shank3 haploinsufficiency is strongly associated with ASD and intellectual disability (Betancur and Buxbaum, 2013). In mouse models, Shank3 loss generates a range of behavior phenotypes from excessive grooming to abnormalities in social behavior (Bey et al., 2018; Bozdagi et al., 2010; Jaramillo et al., 2017; Orefice et al., 2016; Peca et al., 2011; Schmeisser et al., 2012); the phenotype observed likely depends on the knockout strategy used, as there are many alternatively spliced isoforms of Shank3 and different mouse models show different patterns of loss (Jiang and Ehlers, 2013; Monteiro and Feng, 2017; Speed et al., 2015; Wang et al., 2016; Zhou et al., 2016). A Shank3 overexpression mouse model shows a strong hyperactivity phenotype (Han et al., 2013), and in *Drosophila* both overexpression and loss of the only Shank isoform result in synaptic defects (Harris et al., 2016), suggesting that some neuronal properties are quite sensitive to the absolute levels of Shank3 protein. Both the RNAi and the KO mouse model we use here strongly reduce the major neocortical isoforms of Shank3 (Peca et al., 2011; Verpelli et al., 2011), and we find that a ∼50% loss is sufficient to completely abolish homeostatic plasticity *in vitro*. In contrast, overexpression of Shank3 (during rescue of knockdown) had no impact on the ability of neurons to express scaling up, suggesting that too little and too much Shank3 disrupt circuit function through distinct mechanisms.

While an acute, cell-autonomous reduction in Shank3 is sufficient to completely block homeostatic plasticity, this manipulation had little impact on basal synaptic or intrinsic properties. This suggests that acute Shank3 loss disrupts a signaling pathway or pathways critical for the expression of homeostatic adjustments, rather than directly perturbing excitability. The complete signaling pathway through which Shank3 loss disrupts homeostatic plasticity remains to be elucidated, but the ability of Li - and inhibition of Li’s known target GSK3 - to rescue homeostatic plasticity strongly implicates dysregulation of GSK3 in this process. This is consistent with a recent study showing reduced inhibitory phosphorylarion of GSK3 following Shank3 knockdown (Bidinosti et al., 2016). GSK3 hyperactivity targets numerous potential pathways that could contribute to the loss of homeostatic plasticity, among them increased degradation of β-catenin (Orford et al., 1997), a molecule previously implicated in synaptic scaling (Mao et al., 2018; Okuda et al., 2007).

Li is still the first-line treatment for bipolar disorder, and can be efficacious in the treatment of other neuropsychiatric disorders that are unresponsive to standard medications (Rybakowski, 2011), including some human Shankpathies (Egger et al., 2017; Serret et al., 2015). However, despite decades of therapeutic use there is still little understanding of Li’s mode of action. Many studies have investigated the synaptic effect of Li treatment by focusing on NMDA and AMPA receptor function, with complex results that depend on cell type and treatment length. For example, long-term Li treatment downregulates NMDAR function (Nonaka et al., 1998), while shorter treatment at high doses (5 mM) upregulate NMDAR function (Monaco et al., 2018), and 24 hr at a therapeutic concentation has no effect (Sourial-Bassillious et al., 2009). More relevant to the present study is the impact of Li on AMPAR function. Chronic Li treatment decreases synaptic AMPAR levels and reduces AMPA-mediated quantal amplitude (Ankolekar and Sikdar, 2015; Du et al., 2003; Gideons et al., 2017; Gray et al., 2003). In contrast, shorter (16-20 hrs of 2 mM) Li treatment is reported to either increases (Farooq et al., 2017), or decrease (Gray et al., 2003) surface GluA2 levels. Some differences between studies may reflect distinct responses of different neuronal types, as feeding mice Li chow increases AMPAR accumulation in hippocampal but not neocortical neurons (Gould et al., 2008). We find that Li treatment (24 hr at 1 mM) affects neither basal mEPSC nor intrinsic excitability of neocortical pyramidal neurons, but completely rescues the expression of homeostatic plasticity. While it seems likely that chronic Li treatment has additional targets beyond the restoration of homeostatic plasticity, taken together our data suggest that rescue of homeostatic plasticity is one important mechanism of action of Li therapy in human Shankopathies, and possibly in other neuropsychiatric disorders as well.

Synaptic scaling and intrinsic homeostatic plasticity are thought to play essential stabilizing functions during experience-dependent plasticity in sensory systems (Gainey and Feldman, 2017), and although their roles in other brain regions remain largely unexplored, they are widely expressed (Glazewski et al., 2017; Keck et al., 2017). While synaptic scaling defects have been observed in other ASD-associated monogenic disease models such as MeCP2 loss (Blackman et al., 2012, etc), defects in firing rate homeostasis have not previously been examined in any disease model. Here we show that Shank3 loss severely impairs the ability of sensory circuits in visual cortex to undergo firing rate homeostasis and so maintain stable firing in the face of sensory or other perturbations. This is consistent with a large literature documenting alterations in circuit function in Shank3 KO mice, including disturbances in the distribution of firing rates (Peixoto et al., 2016; Wang et al., 2016). Some circuits (striatum, neocortex) appear hyperexcitable (Peixoto et al., 2016), whereas other circuits (hippocampus, cingulate cortex and nucleus accumbens) appear hypoexcitable (Bozdagi et al., 2010; Wang et al., 2016), and a range of synaptic and excitability defects have also been reported across distinct brain regions, cell types, and developmental stages (Jiang and Ehlers, 2013; Kouser et al., 2013; Peixoto et al., 2016; Wang et al., 2017; Zhu et al., 2018). Loss of MeCP2 (another monogenic ASD model) also impairs synaptic scaling (Blackman et al., 2012; Dani et al., 2005; Qiu et al., 2012)and induces a similarly diverse set of changes to excitability in distinct brain regions (Nelson and Valakh, 2015). Our data suggest that some of this diversity in circuit and synaptic outcomes originates from fundamental defects in the homeostatic mechanisms that normally constrain excitability. In the absence of bidirectional homeostatic mechanisms, neurons are at the mercy of perturbations to excitability driven by experience or development, and as these perturbations accumulate over time they will eventually result in dramatic changes in the excitability of particular microcircuits. Furthermore, distinct perturbations to different circuits or cell types would be expected to drive a unique set of pathological changes in each brain region, as observed. Our findings suggest that the loss of key cellular mechanisms for homeostatic compensation is a critical endophenotype that could contribute to a range of circuit disfunctions in Shankopathies.

## Acknowledgments

We thank Dr. Chiara Verpelli for Shank3 SH, Rescue and EV constructs. We thank Lirong Wang, Genelle Rankin and Jorden Rabasco for technical support for culture experiments. We thank Ted Brookings for MATLAB program for analyzing *f*-I data.

## Author contributions

Conceptualization: VT, GT; Methodology/software: VT, GT, ATP, KH, NM; Investigation: VT, NM, ATP, KH, HL; Formal Analysis, VT, ATP, NM, HL; Writing, VT, GT, ATP; Visualization, VT, ATP; Resources, FFW; Supervision/Administration/Funding, GT.

## Data and materials availability

BRD0320was supplied by the Stanley Center for Psychiatric Research, and was confirmed to be of >95% purity based on HPLC LC-MS analysis (Alliance 2795, Waters). All data are presented in main text or supplementary materials, and will be made available upon request.

## Supplemental figure titles

**Fig S1.** Short hairpin expression efficiently reduces Shank3 protein levels.

**Fig S2.** Shank3 KD blocks the activity-dependent changes in synaptic AMPAR accumulation that underlie scaling up and down.

**Fig S3.** Li does not impact magnitude of synaptic scaling.

**Fig S4.** Pooled Passive properties for all conditions.

**Table S1.** Details of statistics.

## STAR METHODS

## CONTACT FOR REAGENT AND RESOURCE SHARING

Further information and requests for resources and reagents should be directed to and will be fulfilled by the Lead Contact, Dr. Gina Turrigiano.

## EXPERIMENTAL MODEL AND SUBJECT DETAILS

For all experiments, mice or rats of both sexes were used. Founder mice were obtained from Jackson labs (Stock No: 017688). Genotyping was done based on primers as previously described (Peca et al., 2011). Long-Evans rats were obtained from Charles River, Wilmington MA. Numbers and ages for each experiment are given below.

## METHOD DETAILS

### Primary cortical neuron culture

Dissociated cultures were prepared from the visual cortex of postnatal day 1 (P1)–P3 Long-Evans rat pups and plated onto a layer of confluent astrocytes on glass-bottom dishes as described previously (Pratt et al., 2003). All experiments were performed after 7-11 days *in vitro* (DIV). All experiments on cultured neurons were performed on neurons with a pyramidal morphology as described (Pratt et al., 2003). All transfections were carried out using Lipofectamine 2000 (Thermo Fisher). Transfected neurons were identified by GFP fluorescence. All control and treatment conditions were run in parallel on sister cultures to minimize variation from individual culture preps and were repeated on a minimum of 3 independent cultures.

### Immunocytochemistry and Image analysis

Neurons were transfected on DIV 6-9 for 2-2.5 days with Lipofectamine 2000, and treated with 5 uM TTX or 100 μM PTX for 24 hrs (unless otherwise specified) prior to fixation in 3.7 % PFA/5% sucrose. Surface GluA2 immunolabeling was performed under non-permeabilizing conditions and all other immunolabeling was performed under permeabilizing conditions with 0.25% Triton X-100. Primary antibodies included Shank3 α gp (Synaptic Systems 162 304), Shank3 α rab (1:400) (162 302Synaptic Systems), Vglut1 (1:500) (catalog no. 135 304, Synaptic Systems), GluA2 (1:1000) α ms (gift from Gouaux lab, OHSU). Immunostained neurons were coverslipped and imaged on a Leica SP5 or Zeiss 880 confocal microscope. Digital images were analyzed in Metamorph (Molecular Devices). For ROI analysis of total intensity, dendritic ROI’s were drawn by hand and background subtracted using an ROI without neurons. Punctate Shank3 or other signals from synaptic markers was identified using the Granularity function in Metamorph. Granules present in both channels were identified as colocalized (a minimum of 3 pixel overlap) and intensities were quantified as previously described (Gainey et al., 2009). Dendritic regions on the apical-like dendrites were analyzed to ensure uniformity in comparisons across neurons.

### Culture Electrophysiology

recordings were performed in whole cell voltage clamp at room temperature; holding potential was −70 mV. Neurons with pyramidal morphology were targeted by visual inspection. Bath solution was ACSF containing (in mM): 126 NaCl, 5.5 KCl, 2 MgSO4, 1 NaH2PO4, 25 NaHCO3, 2 CaCl2, 14 dextrose; and 25 uM picrotoxin, 25 uM D-amino-5-phosphovaleric acid (AP5), and 0.1 μM TTX to isolate AMPA-mediated mEPSCs. Internal solution composition (in mM) was: 120 KMeSO_4_, 10 KCl, 2 MgSO_4_, 10 K-Hepes, 0.5 EGTA, 3 K_2_ATP, 0.3 NaGTP, and 10 Na_2_phosphocreatine. Dextrose was added to adjust osmolarity to 320-330 mOsm. Neurons were excluded if resting membrane potential (Vm) was > −55 mV, series resistance (Rs) was > 20 MΩ, input resistance (Rin) was <100 MΩ, Rin or Vm changed by >30%, or < 25 mEPSCs were obtained. MEPSCs were detected and analyzed using in-house software; detection criteria included amplitudes > 5 pA and rise times < 3 ms. To construct cumulative histograms the first 25 events for each neuron were included. For *f-I* curves transfected neurons were recorded in ACSF with 25 uM Picrotoxin, 50 uM AP5 and 25 uM DNQX. Cells were recorded in whole cell configuration in current clamp and a series of depolarizing step ranging from 30 pA to 250 pA were delivered. Data from EV transfected cells and untransfected control cells were not different and were pooled for some experiments. *f*-I recordings were analyzed using custom software written in house in IGOR pro or MATLAB. All mEPSC and *f*-I data were analyzed blind to condition.

### Slice Electrophysiology

Coronal brain slices containing V1 were prepared as described (Lambo and Turrigiano, 2013). Monocular V1 (V1m) was identified, and whole-cell patch-clamp recordings were obtained from layer 2/3 pyramidal neurons, as previously described (Lambo and Turrigiano, 2013). V1m was identified using the mouse brain atlas (Paxinos and Franklin 2001), using the shape and morphology of the white matter as a reference. Neurons were visualized with a 40x water-immersion objective using infrared-differential interference contrast optics. Internal recording solution contained (in mM) 115 Cs-Methanesulfonate, 10 HEPES(Fisher), 10 BAPTA 4Cs(Fisher), 5.37 Biocytin(Invitrogen), 2 QX-314 Cl(Tocris), 1.5 MgCl2, 1 EGTA, 10 Na2-Phosphocreatine, 4 ATP-Mg, 0.3 GTP-Na, with sucrose added to bring osmolality to 295, and CsOH added to bring pH to 7.35. For mEPSC recordings, neurons were voltage clamped to −70 mV in standard ACSF containing TTX (0.2 uM), APV (50 uM), and picrotoxin (25 uM), continuously oxygenated and warmed to 34^°^C. Neurons were excluded from analysis if series resistance was > 20 MΩ or if input resistance was < 80 MΩ. Pyramidal neurons were identified and targeted by the presence of an apical dendrite and tear-drop shaped soma, and morphology was confirmed by *post hoc* reconstruction of biocytin fills.

### Reagents

GSK3i (BRD0320-Broad Institute), LiCl2 (Sigma), TTX (Tocris), PTX (Sigma), DNQX (Sigma), AP5 (Tocris), DMSO (Sigma), Alexa 488, 555, 647 (Invitrogen).

### DNA constructs

Shank3 SH and Rescue plasmids were obtained from Dr. Chiara Verpelli and were previously characterized.(Verpelli et al., 2011)

### Surgery and *in vivo* recordings

Electrode array implantations were carried out on littermate Shank3 and WT mice generated from Het-Het matings (WT: n = 7, KO: n = 7; P21-24 at time of surgery). Mice were implanted unilaterally under isofluorane anesthesia with custom 16-channel 33 μm tungsten microelectrode arrays (Tucker-Davis Technologies, Alachua, FL, wire diameter 33 μm, electrode spacing 250 μm, row separation 375 μm, tip angle 45^°^) into V1. Wires were cut to different lengths so as to span all layers of V1. Surgical procedure was as described previously (Hengen et al., 2013). Briefly, the skull was exposed and cleaned and two holes were drilled above the cerebellum and right hemisphere to support stainless steel miniature screws. After drilling a craniotomy, the electrode array was inserted in V1 in the left hemisphere to a depth of 1.0 mm for the longest wire and grounded to both screws. Dental cement was used to connect the array to the entirety of the exposed bone surface as well as the steel screws. The total headcap weight was less than 0.8 grams.

Animals were allowed to recover for two full days post-surgery in transparent plastic cages with *ad libitum* access to food and water. Recording began on the third day after surgery. Animals were allowed to habituate to the recording chamber for ∼12 hours prior to the start of the experiment. The recording chamber (12”x12”, plexiglass) was lined with 1.5” of bedding and housed two littermates (one Shank3 KO and one WT mouse). Animals had *ad libitum* food and water and were separated by a clear plastic divider with several holes allowing for tactile and olfactory interaction but preventing jostling of headcaps and electrode arrays. Novel toys were introduced every 24h, to enrich the animals’ experience and promote activity and exploration. Lighting and temperature were kept constant (LD 12:12, lights on at 7:30 am, 21^°^ C, humidity 25%-55%).

Data were collected continuously for nine days (∼200 hours). After lights off on the third day of baseline recording animals were removed from the recording chamber and placed under isoflurane anesthesia. The eyelid contralateral to the implanted hemisphere was sutured. The procedure lasted ∼15 min per animal. Mice were fully recovered from anesthesia 15 minutes after re-introduction in the recording chamber. The lid suture was kept in place for 6 days. Electrode placement was verified histologically at the end of each experiment.

### *In vivo* data analysis

Voltage signals were collected at 25 kHz and streamed to disk using a Tucker-Davis Technologies Neurophysiology Workstation and Data Streamer. Spike extraction was performed offline using custom software (MATLAB). Briefly, a 500 kHz high-pass filter was applied and threshold crossings (threshold: −4 standard deviations from the mean signal) were detected and saved (8 samples before and 24 samples after the crossing). Spike waveforms were re-sampled at 3x the original sampling rate, then subjected to principal component analysis (PCA) and the first four principal components were used to cluster each channel’s waveforms using the KlustaKwik algorithm (Harris et al., 2000). Clusters were not cut manually; instead, they were merged or trimmed as described previously (Hengen et al., 2016).

Spike sorting was performed using custom software (MATLAB) that relies on a machine learning approach. For each cluster, a set of 19 features was extracted, including ISI contamination (percentage of ISIs < 3 ms), similarity to regular spiking unit (RSU) and fast-spiking (FS) waveform templates, 60 Hz noise contamination, rise and decay time and slope of the mean waveform, and waveform amplitude and width. Cluster quality was also ensured by thresholding of L-Ratio and Mahalanobis distance (Schmitzer-Torbert et al., 2005). The set of resulting features was fed to a random forest classifier, trained on a set of 1200 clusters scored by human experts (ATP and KBH). Each cluster was assigned a quality value between 1 and 4, with 1 and 2 representing single-unit data. Each cluster also had a confidence score for its classification, ranging from 0% to 100%. Only units with quality 1 or 2 were used for analysis. Clusters with quality 2 and a confidence score less than 70% were excluded. Only regular spiking (putative pyramidal) units (RSUs) were used for analysis. Cells were classified into RSUs or fast spiking units (FSUs) based on the following mean waveform parameters: time between the negative peak and the first subsequent positive peak (trough-to-peak time); slope of the waveform from 0.4 msec after the negative peak to last sample (tail slope). RSUs were identified as having a tail slope > 0.005 and trough-to-peak time > 0.39 msec (Niell and Stryker, 2008).

Recordings were time-stamped so that time 0 represents 7:30 am (lights on) on the first baseline day. The recording was split into nine 24-hour periods starting at this time. The set of spikes from each period was fed to the clustering and spike-sorting algorithm. The majority of cells could not be followed continuously over these 9 day experiments, so instead we compared the ensemble averages and distributions for all RSUs detected for each day. The criteria used to determine when a cell was on- or off-line were as described previously(Hengen et al., 2016); to obtain reasonable estimates of mean firing rates only neurons recorded for >2 hours were included.

### Histology

At the end of each experiment, animals were administered a lethal dose of ketamine/xylazine (dosage: 200 mg/kg and 7 mg/kg respectively) and transcardially perfused with 0.9% saline followed by 4% paraformaldehyde. Brains were postfixed in 10% formalin and 30% sucrose solution for >3 days. Brains were then sectioned at 60 μm, mounted on glass slides, stained using cresyl violet according to a standard Nissl staining protocol, and imaged using a light microscope. Electrode tracks were visually identified in stained sections. Data from electrodes that were not in V1 were excluded from analyses.

## QUANTIFICATION AND STATISTICAL ANALYSIS

The statistical test(s) used for each dataset are indicated in the figure legends. For comparison of two groups two tailed t-test were used as described. Comparison among multiple groups was via one-way ANOVA in conjunction with Levene’s test for homogeneity. This was followed by post hoc tests to adjust for multiple comparisons with Bonferroni’s correction as noted. Comparison of multiple non-normally distributed groups was via the Kruskal-Wallis test, followed by pairwise Wilcoxon rank-sum tests adjusted for multiple comparisons using the Benjamini-Hochberg procedure (false discovery rate = 0.05). Comparison of cumulative distributions was done with Kruskal–Wallis test or Anderson-Darling test followed by either Bonferroni’s correction or Benjamini-Hochberg FDR procedure (FDR = 0.05). P-values below 0.05 were considered significant. The actual P values of each comparison conducted are given in Supplementary Table 1.

## References and Notes

Abbott, L.F., and Nelson, S.B. (2000). Synaptic plasticity: taming the beast. Nat Neurosci 3 Suppl, 1178–1183.

Ankolekar, S.M., and Sikdar, S.K. (2015). Early postnatal exposure to lithium in vitro induces changes in AMPAR mEPSCs and vesicular recycling at hippocampal glutamatergic synapses. J Biosci 40, 339–354.

Betancur, C., and Buxbaum, J.D. (2013). SHANK3 haploinsufficiency: a “common” but underdiagnosed highly penetrant monogenic cause of autism spectrum disorders. Mol Autism 4, 17.

Bey, A.L., Wang, X., Yan, H., Kim, N., Passman, R.L., Yang, Y., Cao, X., Towers, A.J., Hulbert, S.W., Duffney, L.J., et al. (2018). Brain region-specific disruption of Shank3 in mice reveals a dissociation for cortical and striatal circuits in autism-related behaviors. Transl Psychiatry 8, 94.

Bidinosti, M., Botta, P., Kruttner, S., Proenca, C.C., Stoehr, N., Bernhard, M., Fruh, I., Mueller, M., Bonenfant, D., Voshol, H., et al. (2016). CLK2 inhibition ameliorates autistic features associated with SHANK3 deficiency. Science 351, 1199–1203.

Blackman, M.P., Djukic, B., Nelson, S.B., and Turrigiano, G.G. (2012). A critical and cell-autonomous role for MeCP2 in synaptic scaling up. J Neurosci 32, 13529–13536.

Bozdagi, O., Sakurai, T., Papapetrou, D., Wang, X., Dickstein, D.L., Takahashi, N., Kajiwara, Y., Yang, M., Katz, A.M., Scattoni, M.L., et al. (2010). Haploinsufficiency of the autism-associated Shank3 gene leads to deficits in synaptic function, social interaction, and social communication. Mol Autism 1, 15.

Dani, V.S., Chang, Q., Maffei, A., Turrigiano, G.G., Jaenisch, R., and Nelson, S.B. (2005). Reduced cortical activity due to a shift in the balance between excitation and inhibition in a mouse model of Rett syndrome. Proc Natl Acad Sci U S A 102, 12560–12565.

Darville, H., Poulet, A., Rodet-Amsellem, F., Chatrousse, L., Pernelle, J., Boissart, C., Heron, D., Nava, C., Perrier, A., Jarrige, M., et al. (2016). Human Pluripotent Stem Cell-derived Cortical Neurons for High Throughput Medication Screening in Autism: A Proof of Concept Study in SHANK3 Haploinsufficiency Syndrome. EBioMedicine 9, 293–305.

Davis, G.W. (2013). Homeostatic signaling and the stabilization of neural function. Neuron 80, 718–728.

Desai, N.S., Rutherford, L.C., and Turrigiano, G.G. (1999). Plasticity in the intrinsic excitability of cortical pyramidal neurons. Nat Neurosci 2, 515–520.

Du, J., Gray, N.A., Falke, C., Yuan, P., Szabo, S., and Manji, H.K. (2003). Structurally dissimilar antimanic agents modulate synaptic plasticity by regulating AMPA glutamate receptor subunit GluR1 synaptic expression. Ann N Y Acad Sci 1003, 378–380.

Egger, J.I.M., Verhoeven, W.M.A., Groenendijk-Reijenga, R., and Kant, S.G. (2017). Phelan-McDermid syndrome due to SHANK3 mutation in an intellectually disabled adult male: successful treatment with lithium. BMJ Case Rep 2017.

Farooq, M., Kim, S., Patel, S., Khatri, L., Hikima, T., Rice, M.E., and Ziff, E.B. (2017). Lithium increases synaptic GluA2 in hippocampal neurons by elevating the delta-catenin protein. Neuropharmacology 113, 426–433.

Freland, L., and Beaulieu, J.M. (2012). Inhibition of GSK3 by lithium, from single molecules to signaling networks. Front Mol Neurosci 5, 14.

Gainey, M.A., and Feldman, D.E. (2017). Multiple shared mechanisms for homeostatic plasticity in rodent somatosensory and visual cortex. Philos Trans R Soc Lond B Biol Sci 372.

Gainey, M.A., Hurvitz-Wolff, J.R., Lambo, M.E., and Turrigiano, G.G. (2009). Synaptic scaling requires the GluR2 subunit of the AMPA receptor. J Neurosci 29, 6479–6489.

Gideons, E.S., Lin, P.Y., Mahgoub, M., Kavalali, E.T., and Monteggia, L.M. (2017). Chronic lithium treatment elicits its antimanic effects via BDNF-TrkB dependent synaptic downscaling. Elife 6.

Glazewski, S., Greenhill, S., and Fox, K. (2017). Time-course and mechanisms of homeostatic plasticity in layers 2/3 and 5 of the barrel cortex. Philos Trans R Soc Lond B Biol Sci 372.

Gray, N.A., Zhou, R., Du, J., Moore, G.J., and Manji, H.K. (2003). The use of mood stabilizers as plasticity enhancers in the treatment of neuropsychiatric disorders. J Clin Psychiatry 64 Suppl 5, 3–17.

Han, K., Holder, J.L., Jr., Schaaf, C.P., Lu, H., Chen, H., Kang, H., Tang, J., Wu, Z., Hao, S., Cheung, S.W., et al. (2013). SHANK3 overexpression causes manic-like behaviour with unique pharmacogenetic properties. Nature 503, 72–77.

Harris, K.D., Henze, D.A., Csicsvari, J., Hirase, H., and Buzsaki, G. (2000). Accuracy of tetrode spike separation as determined by simultaneous intracellular and extracellular measurements. J Neurophysiol 84, 401–414.

Harris, K.P., Akbergenova, Y., Cho, R.W., Baas-Thomas, M.S., and Littleton, J.T. (2016). Shank Modulates Postsynaptic Wnt Signaling to Regulate Synaptic Development. J Neurosci 36, 5820–5832.

Hengen, K.B., Lambo, M.E., Van Hooser, S.D., Katz, D.B., and Turrigiano, G.G. (2013). Firing rate homeostasis in visual cortex of freely behaving rodents. Neuron 80, 335–342.

Hengen, K.B., Torrado Pacheco, A., McGregor, J.N., Van Hooser, S.D., and Turrigiano, G.G. (2016). Neuronal Firing Rate Homeostasis Is Inhibited by Sleep and Promoted by Wake. Cell 165, 180–191.

Jaramillo, T.C., Speed, H.E., Xuan, Z., Reimers, J.M., Escamilla, C.O., Weaver, T.P., Liu, S., Filonova, I., and Powell, C.M. (2017). Novel Shank3 mutant exhibits behaviors with face validity for autism and altered striatal and hippocampal function. Autism Res 10, 42–65.

Jiang, Y.H., and Ehlers, M.D. (2013). Modeling autism by SHANK gene mutations in mice. Neuron 78, 8–27.

Joseph, A., and Turrigiano, G.G. (2017). All for One But Not One for All: Excitatory Synaptic Scaling and Intrinsic Excitability Are Coregulated by CaMKIV, Whereas Inhibitory Synaptic Scaling Is Under Independent Control. J Neurosci 37, 6778–6785.

Kaneko, M., Stellwagen, D., Malenka, R.C., and Stryker, M.P. (2008). Tumor necrosis factor-alpha mediates one component of competitive, experience-dependent plasticity in developing visual cortex. Neuron 58, 673–680.

Keck, T., Keller, G.B., Jacobsen, R.I., Eysel, U.T., Bonhoeffer, T., and Hubener, M. (2013). Synaptic scaling and homeostatic plasticity in the mouse visual cortex in vivo. Neuron 80, 327–334.

Keck, T., Toyoizumi, T., Chen, L., Doiron, B., Feldman, D.E., Fox, K., Gerstner, W., Haydon, P.G., Hubener, M., Lee, H.K., et al. (2017). Integrating Hebbian and homeostatic plasticity: the current state of the field and future research directions. Philos Trans R Soc Lond B Biol Sci 372.

Kouser, M., Speed, H.E., Dewey, C.M., Reimers, J.M., Widman, A.J., Gupta, N., Liu, S., Jaramillo, T.C., Bangash, M., Xiao, B., et al. (2013). Loss of predominant Shank3 isoforms results in hippocampus-dependent impairments in behavior and synaptic transmission. J Neurosci 33, 18448–18468.

Lambo, M.E., and Turrigiano, G.G. (2013). Synaptic and intrinsic homeostatic mechanisms cooperate to increase L2/3 pyramidal neuron excitability during a late phase of critical period plasticity. J Neurosci 33, 8810–8819.

Mao, W., Salzberg, A.C., Uchigashima, M., Hasegawa, Y., Hock, H., Watanabe, M., Akbarian, S., Kawasawa, Y.I., and Futai, K. (2018). Activity-Induced Regulation of Synaptic Strength through the Chromatin Reader L3mbtl1. Cell Rep 23, 3209–3222.

Marder, E., and Goaillard, J.M. (2006). Variability, compensation and homeostasis in neuron and network function. Nat Rev Neurosci 7, 563–574.

Monaco, S.A., Ferguson, B.R., and Gao, W.J. (2018). Lithium Inhibits GSK3beta and Augments GluN2A Receptor Expression in the Prefrontal Cortex. Front Cell Neurosci 12, 16.

Monteiro, P., and Feng, G. (2017). SHANK proteins: roles at the synapse and in autism spectrum disorder. Nat Rev Neurosci 18, 147–157.

Mrsic-Flogel, T.D., Hofer, S.B., Ohki, K., Reid, R.C., Bonhoeffer, T., and Hubener, M. (2007). Homeostatic regulation of eye-specific responses in visual cortex during ocular dominance plasticity. Neuron 54, 961–972.

Nelson, S.B., and Valakh, V. (2015). Excitatory/Inhibitory Balance and Circuit Homeostasis in Autism Spectrum Disorders. Neuron 87, 684–698.

Niell, C.M., and Stryker, M.P. (2008). Highly selective receptive fields in mouse visual cortex. J Neurosci 28, 7520–7536.

Nonaka, S., Hough, C.J., and Chuang, D.M. (1998). Chronic lithium treatment robustly protects neurons in the central nervous system against excitotoxicity by inhibiting N-methyl-D-aspartate receptor-mediated calcium influx. Proc Natl Acad Sci U S A 95, 2642–2647.

Okuda, T., Yu, L.M., Cingolani, L.A., Kemler, R., and Goda, Y. (2007). beta-Catenin regulates excitatory postsynaptic strength at hippocampal synapses. Proc Natl Acad Sci U S A 104, 13479–13484.

Orefice, L.L., Zimmerman, A.L., Chirila, A.M., Sleboda, S.J., Head, J.P., and Ginty, D.D. (2016). Peripheral Mechanosensory Neuron Dysfunction Underlies Tactile and Behavioral Deficits in Mouse Models of ASDs. Cell 166, 299–313.

Orford, K., Crockett, C., Jensen, J.P., Weissman, A.M., and Byers, S.W. (1997). Serine phosphorylation-regulated ubiquitination and degradation of beta-catenin. J Biol Chem 272, 24735–24738.

Peca, J., Feliciano, C., Ting, J.T., Wang, W., Wells, M.F., Venkatraman, T.N., Lascola, C.D., Fu, Z., and Feng, G. (2011). Shank3 mutant mice display autistic-like behaviours and striatal dysfunction. Nature 472, 437–442.

Peixoto, R.T., Wang, W., Croney, D.M., Kozorovitskiy, Y., and Sabatini, B.L. (2016). Early hyperactivity and precocious maturation of corticostriatal circuits in Shank3B(-/-) mice. Nat Neurosci 19, 716–724.

Pisanu, C., Melis, C., and Squassina, A. (2016). Lithium Pharmacogenetics: Where Do We Stand? Drug Dev Res 77, 368–373.

Pratt, K.G., Watt, A.J., Griffith, L.C., Nelson, S.B., and Turrigiano, G.G. (2003). Activity-dependent remodeling of presynaptic inputs by postsynaptic expression of activated CaMKII. Neuron 39, 269–281.

Qiu, Z., Sylwestrak, E.L., Lieberman, D.N., Zhang, Y., Liu, X.Y., and Ghosh, A. (2012). The Rett syndrome protein MeCP2 regulates synaptic scaling. J Neurosci 32, 989–994.

Rybakowski, J.K. (2011). Lithium in neuropsychiatry: a 2010 update. World J Biol Psychiatry 12, 340–348.

Schmeisser, M.J., Ey, E., Wegener, S., Bockmann, J., Stempel, A.V., Kuebler, A., Janssen, A.L., Udvardi, P.T., Shiban, E., Spilker, C., et al. (2012). Autistic-like behaviours and hyperactivity in mice lacking ProSAP1/Shank2. Nature 486, 256–260.

Serret, S., Thummler, S., Dor, E., Vesperini, S., Santos, A., and Askenazy, F. (2015). Lithium as a rescue therapy for regression and catatonia features in two SHANK3 patients with autism spectrum disorder: case reports. BMC Psychiatry 15, 107.

Smith, G.B., Heynen, A.J., and Bear, M.F. (2009). Bidirectional synaptic mechanisms of ocular dominance plasticity in visual cortex. Philos Trans R Soc Lond B Biol Sci 364, 357–367.

Sourial-Bassillious, N., Rydelius, P.A., Aperia, A., and Aizman, O. (2009). Glutamate-mediated calcium signaling: a potential target for lithium action. Neuroscience 161, 1126–1134.

Speed, H.E., Kouser, M., Xuan, Z., Reimers, J.M., Ochoa, C.F., Gupta, N., Liu, S., and Powell, C.M. (2015). Autism-Associated Insertion Mutation (InsG) of Shank3 Exon 21 Causes Impaired Synaptic Transmission and Behavioral Deficits. J Neurosci 35, 9648–9665.

Turrigiano, G.G. (2017). The dialectic of Hebb and homeostasis. Philos Trans R Soc Lond B Biol Sci 372.

Turrigiano, G.G., Leslie, K.R., Desai, N.S., Rutherford, L.C., and Nelson, S.B. (1998). Activity-dependent scaling of quantal amplitude in neocortical neurons. Nature 391, 892–896.

Turrigiano, G.G., and Nelson, S.B. (2004). Homeostatic plasticity in the developing nervous system. Nat Rev Neurosci 5, 97–107.

Verpelli, C., Dvoretskova, E., Vicidomini, C., Rossi, F., Chiappalone, M., Schoen, M., Di Stefano, B., Mantegazza, R., Broccoli, V., Bockers, T.M., et al. (2011). Importance of Shank3 protein in regulating metabotropic glutamate receptor 5 (mGluR5) expression and signaling at synapses. J Biol Chem 286, 34839–34850.

Wagner, F.F., Bishop, J.A., Gale, J.P., Shi, X., Walk, M., Ketterman, J., Patnaik, D., Barker, D., Walpita, D., Campbell, A.J., et al. (2016). Inhibitors of Glycogen Synthase Kinase 3 with Exquisite Kinome-Wide Selectivity and Their Functional Effects. ACS Chem Biol 11, 1952–1963.

Wang, W., Li, C., Chen, Q., van der Goes, M.S., Hawrot, J., Yao, A.Y., Gao, X., Lu, C., Zang, Y., Zhang, Q., et al. (2017). Striatopallidal dysfunction underlies repetitive behavior in Shank3-deficient model of autism. J Clin Invest 127, 1978–1990.

Wang, X., Bey, A.L., Katz, B.M., Badea, A., Kim, N., David, L.K., Duffney, L.J., Kumar, S., Mague, S.D., Hulbert, S.W., et al. (2016). Altered mGluR5-Homer scaffolds and corticostriatal connectivity in a Shank3 complete knockout model of autism. Nat Commun 7, 11459.

Wilson, H.L., Wong, A.C., Shaw, S.R., Tse, W.Y., Stapleton, G.A., Phelan, M.C., Hu, S., Marshall, J., and McDermid, H.E. (2003). Molecular characterisation of the 22q13 deletion syndrome supports the role of haploinsufficiency of SHANK3/PROSAP2 in the major neurological symptoms. J Med Genet 40, 575–584.

Yi, F., Danko, T., Botelho, S.C., Patzke, C., Pak, C., Wernig, M., and Sudhof, T.C. (2016). Autism-associated SHANK3 haploinsufficiency causes Ih channelopathy in human neurons. Science 352, aaf2669.

Zhou, Y., Kaiser, T., Monteiro, P., Zhang, X., Van der Goes, M.S., Wang, D., Barak, B., Zeng, M., Li, C., Lu, C., et al. (2016). Mice with Shank3 Mutations Associated with ASD and Schizophrenia Display Both Shared and Distinct Defects. Neuron 89, 147–162.

Zhu, M., Idikuda, V.K., Wang, J., Wei, F., Kumar, V., Shah, N., Waite, C.B., Liu, Q., and Zhou, L. (2018). Shank3-deficient thalamocortical neurons show HCN channelopathy and alterations in intrinsic electrical properties. J Physiol 596, 1259–1276.

